# Esophageal epithelial cell-state transitions underlie the severity of pediatric eosinophilic esophagitis

**DOI:** 10.64898/2026.03.07.710295

**Authors:** Yu Wang, Manish Kumar Sinha, Peter Ghattas, Jennifer M Pilat, Yash A Choksi, Hee-Woong Lim, Marc E Rothenberg, Quanhu Sheng, Girish Hiremath, Seesandra V Rajagopala

## Abstract

Eosinophilic esophagitis (EoE) is a leading cause of chronic esophageal dysfunction driven by immune-mediated inflammation. Peak eosinophil count (PEC) in esophageal biopsies is routinely used to assess disease activity, and its associated molecular mechanisms have been well studied. However, PEC only partially captures overall disease severity, which is comprehensively captured by the Index of Severity for Eosinophilic Esophagitis (I-SEE). In contrast to PEC, the molecular and cellular programs associated with the I-SEE-defined disease severity, particularly in children, remain poorly understood. We integrated bulk transcriptomic profiling of pediatric esophageal biopsies with clinical severity metrics and a matched single-cell transcriptomic reference. Increasing severity was associated with a shift from type 2 inflammatory activation toward epithelial stress, cytoskeletal and junctional disruption, metabolic dysfunction, and extracellular matrix remodeling. Single-cell–informed analyses identified that proliferating and transitional epithelial cell states were strongly associated with higher I-SEE scores and exhibited impaired differentiation, heightened metabolic and oxidative stress responses, and structural remodeling programs not captured by bulk transcriptomic analyses alone. These findings reposition epithelial remodeling, rather than eosinophil burden alone, as a central molecular correlate of disease severity in pediatric EoE and provide a framework for improved disease stratification and therapeutic intervention.

## Introduction

Eosinophilic esophagitis (EoE) is a chronic, immune-mediated inflammatory disease of the esophagus. As a clinicopathologic disease, it is a leading cause of esophageal dysfunction across all age^1^. Children with EoE present with feeding difficulties, vomiting, abdominal pain, and growth faltering, symptoms suggestive of an inflammatory phenotype. In contrast, adolescents and young adults present with dysphagia and esophageal food impaction, concerning a fibrostenotic phenotype. An intense esophageal eosinophilia with progressive tissue remodeling, as confirmed histologically, supports the diagnosis of EoE^2^. Type 2 (Th2) inflammatory signaling, exemplified by key effector cytokines and downstream target genes such as interleukin (IL)-4, IL-13, CCL26, POSTN, CAPN14, and the lineage-defining transcription factor GATA3, has been implicated in epithelial injury, subepithelial remodeling, and chronicity^3–8,3–8^(3-8)^3–8^. Additionally, evidence from single-cell RNA (scRNA) sequencing has revealed striking epithelial and immune heterogeneity in EoE, identifying discrete epithelial lineages and immune-epithelial interactions that shape the course of disease^5–8^.

While molecular studies have advanced our understanding of EoE pathogenesis, they have not been fully integrated with clinical measures of the disease burden. At present, peak eosinophil count (PEC) in the esophageal biopsies remains central to diagnosis, disease activity assessment, and management^9^. However, PEC inadequately captures the disease’s broader impact. Accordingly, the Index of Severity for Eosinophilic Esophagitis (I-SEE), a multidimensional framework, was developed to comprehensively assess disease burden. I-SEE integrates patient-reported symptoms, endoscopic findings, histologic features, and complications into a composite severity score. I-SEE categorizes patients into inactive, mild, moderate, and severe disease states, enabling a more comprehensive and clinically meaningful assessment of EoE severity for clinical care and longitudinal monitoring^10^. Sato et al. recently reported that the inactive, mild, and moderate I-SEE scores correlated with molecular features, and pro-melanin-concentrating hormone (PMCH) and endonuclease (ENDOU) were the most positively and negatively correlated markers of disease severity, respectively. However, this secondary analysis used a pre-defined gene panel, which limited discovery, and included a cohort of adults and children^11^. As such, pediatric-specific molecular and cellular correlates of I-SEE have yet to be defined through an unbiased approach.

To address this knowledge gap, we performed bulk RNA sequencing of esophageal biopsies collected from children with and without EoE, integrating these data with comprehensive clinical phenotyping, I-SEE scores, and a matched scRNA-seq reference. Through differential gene expression, pathway enrichment, cell-type deconvolution, and phenotype-guided Scissor analysis, we identified how immune activation, epithelial stress response, and epithelial lineage transitions shape the I-SEE score in children with EoE. Our findings confirm a transcriptomic spectrum that links Th2 inflammation with epithelial stress and fibrotic remodeling, identify subsets of proliferating and transitional epithelial populations as the dominant cellular correlates of severe disease, and uncover severity-specific pathways not captured by eosinophil counts alone. These results provide a molecular interpretation of the I-SEE score in children and establish a foundation for precision stratification and targeted therapeutic development in EoE.

## Results

### Clinical characteristics

We performed bulk RNA sequencing on the esophageal biopsies collected from 121 children [EoE = 75, and control = 46] aged 6-18 years. Based on the conventional threshold, children with ≥15 eosinophils per high-power field (eos/hpf) were classified as active EoE (aEoE, n = 43) and those with < 15 eos/hpf as inactive EoE (iEoE, n = 32)^9^. Demographic characteristics, such as age and ethnicity, were comparable across groups, although a significantly higher proportion of females (p < 0.01) were in the control group. As expected, a significantly higher proportion of children in the EoE group had a history of esophageal food impaction and dilation, and atopic co-morbidities. Children with aEoE had the highest EREFS score [median (range): 2 (0-5)] and PEC [2 (18-112)]. The median [interquartile range (IQR)] I-SEE score was 4 (1.5-5), and 20% were categorized as inactive, 61% as mild, 10% as moderate, and 4% as severe (Table S1). No significant association was observed between symptom duration and the I-SEE score (Supplementary Fig. 1I). One patient had an unexpectedly high I-SEE score, which was driven by malnutrition and endoscopic abnormalities despite being classified as iEoE.

### Transcriptomic distinctions across pediatric EoE activity

First, to delineate how EoE activity (defined by the PEC) affects the esophageal transcriptome, we assessed global gene expression patterns in aEoE, iEoE, and controls. Principal component analysis of the whole transcriptomes showed a clear separation between aEoE and controls. In contrast, iEoE samples clustered closer to the controls (Supplemental Fig. 1A), which was not explained by clinical or technical covariates, including age, sex, treatment, or batch (data not shown). Differential expression analyses demonstrated substantial transcriptional changes in aEoE. The aEoE transcriptomic signature was more coherent and showed stronger effect sizes than the overall EoE group (combining aEoE and iEoE) (Fig. 1A, B). In total, 1,446 differentially expressed genes (DEGs) were identified in aEoE versus control (Fig. 1A), and 435 DEGs were identified in aEoE versus iEoE (Fig. 1C; Supplemental Fig. 1E; Tables S2–S4). In contrast, the overall EoE group showed fewer DEGs (424 vs. control, Fig. 1B), of which 96% overlapped with those identified in aEoE versus control. Notably, iEoE exhibited only a single DEG compared to controls [Regenerating family member 1 alpha (REG1A)]. Furthermore, log2 fold changes (log_2_FC) across all genes were highly correlated between aEoE vs. control and EoE vs. control comparisons (*R* = 0.96, p < 2.2e−16, Supplemental Fig. 1B), indicating that active disease accounts for the majority of the EoE transcriptomic signature. Similarly, log_2_FC values were strongly correlated between aEoE versus control and aEoE versus iEoE (*R* = 0.84, p < 2.2 × 10⁻¹; Supplemental Fig. 1C), indicating minor transcriptional differences in iEoE, consistent with a largely quiescent or partially resolved disease state.

**Fig. 1.**
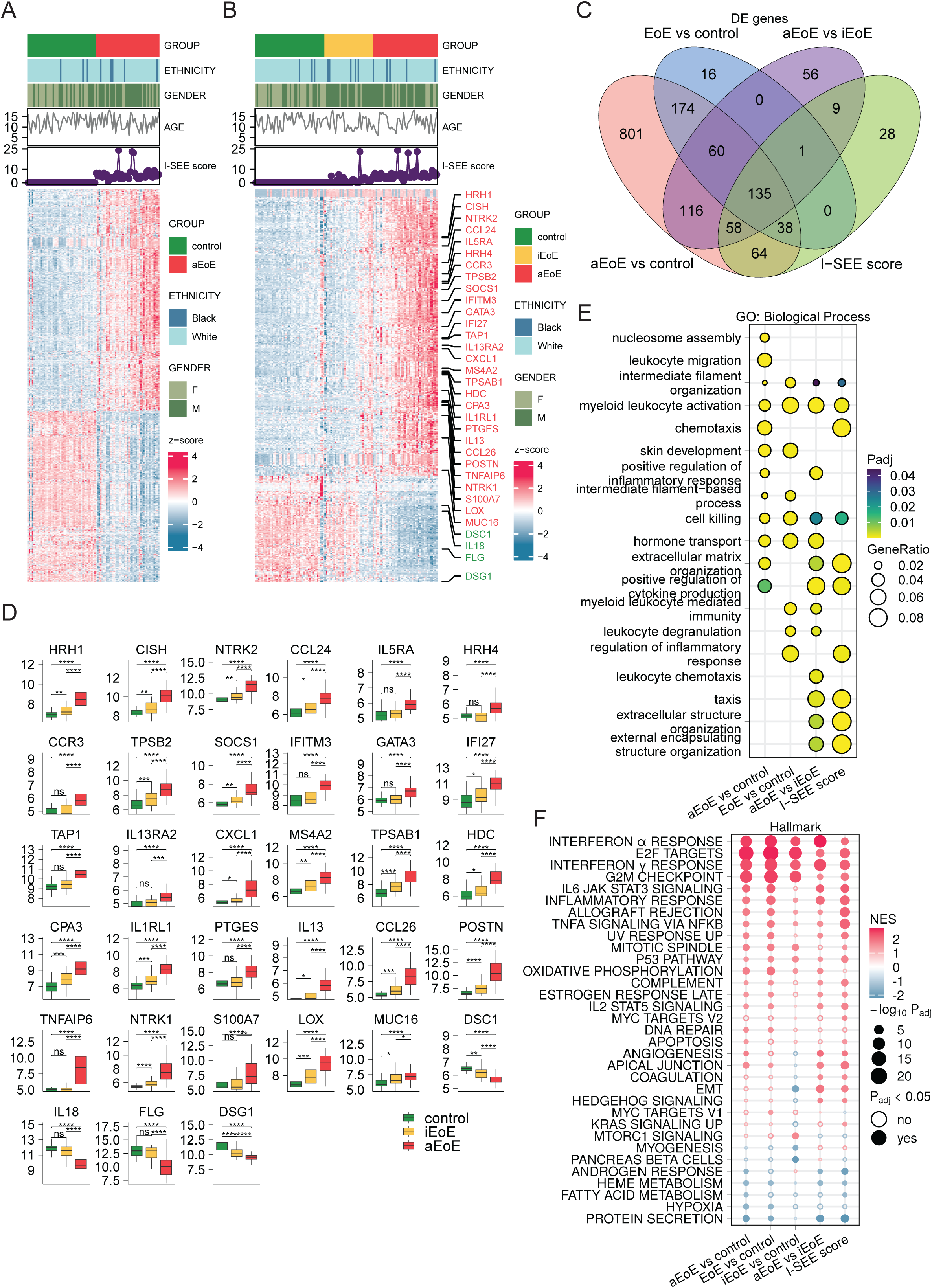
A) Differential expression genes identified by comparing aEoE versus control. Data batch, ethnicity, gender, and age were adjusted as covariates. Red indicates up-regulation in aEoE, while blue indicates up-regulation in control. B) Differential expression genes identified by comparing EoE versus control. C) Venn diagram of DE genes from different comparisons: aEoE vs. control, EoE vs. control, aEoE vs. iEoE, and severity per I-SEE score. D) Boxplots of selected genes to show their expression distribution across groups. The y-axis shows the expression values after normalization with the variance-stabilizing transformation function from DESeq2. The Wilcoxon rank-sum test was applied to assess differences between groups. Multiple test adjustments were performed using the Benjamini-Hochberg procedure. ns: non-significant; *: adjusted P value < 0.05; **: adjusted P value < 0.01; ***: adjusted P value < 001; ****: adjusted P value < 0.0001. E) Top 8 significant GO Biological Process terms of each comparison from function enrichment analysis. Redundant terms were filtered with simplify function (cutoff = 0.6, by = “p.adjust”, select_fun = min, measure = “Wang”). F) Significant Hallmark gene sets from at least one comparison from pre-ranked GSEA analysis. Color indicates the normalized enrichment score, while point size indicates the adjusted p-value (-log10Padj). Up-regulated pathways in the numerator group were shown in red, while the down-regulated pathways in the numerator group were shown in blue.

Transcript-level analysis of representative EoE-related pathways and their associated DEGs (Fig. 1B) illustrated a graded change from control to iEoE to aEoE. Genes involved in type 2 inflammation and eosinophil recruitment (IL13, CCL26, CCL24, IL5RA, IL1RL1, GATA3), mast-cell activation (CPA3, HDC, TPSAB1, TPSB2, MS4A2), neuro-immune signaling (HRH1, HRH4, NTRK1, NTRK2), innate immune activation (CXCL1, IFI27, IFITM3, SOCS1), and tissue remodeling (LOX, POSTN, TNFAIP6) showed graded upregulation from control to iEoE to aEoE, while epithelial barrier structure (FLG, DSG1, DSC1) and IL18 showed graded downregulation from control to iEoE to aEoE (Fig. 1D). IL-18 is an inflammasome-dependent cytokine that amplifies innate inflammation and strongly enhances IFN-γ–mediated Th1 immunity, while also influencing epithelial barrier responses and, under certain cytokine environments, promoting Th2-like effector programs. This coordinated pattern reinforces that iEoE retains partial molecular activity but lacks the full inflammatory and remodeling signatures characteristic of active disease. Collectively, these results highlight the transcriptomic distinctions between the EoE activity states and that combining iEoE and aEoE underestimates the full molecular signature of active disease. Analyzing them separately provides a greater resolution of disease-associated molecular features.

### EoE activity reflects immune activation, epithelial stress, and matrix remodeling

Having demonstrated that transcriptomic alterations scale with the disease activity from control to iEoE to aEoE, we next examined the biological pathways and gene sets that drive molecular escalation.

Gene Ontology (GO) enrichment analysis revealed prominent enrichment for dysregulated immune response, cytokine and interleukin signaling, chromatin and nucleosome organization, extracellular matrix (ECM) organization, epithelial barrier regulation, and leukocyte chemotaxis (Fig. 1E; Tables S6–S8) in EoE compared to controls. These GO-level patterns indicate large-scale disturbances in immune activation, epithelial homeostasis, and tissue structural remodeling in disease development.

Hallmark gene set enrichment analysis further demonstrated that coordinated signaling programs track with disease activity (Fig. 1F; Tables S9–S11). Compared with controls, both iEoE and aEoE showed enrichment of cell cycle activity (E2F targets, G2M checkpoint), interferon responses, inflammatory programs (TNFA/NF-κB), and tissue repair signatures. These findings indicate that, despite the detection of only a single significantly differentially expressed gene in iEoE versus control, genes in these programs are regulated in iEoE in the same direction as observed in aEoE. Meanwhile, the activity-related distinctions emerged when comparing iEoE and aEoE. aEoE displayed suppression of metabolic pathways, including oxidative phosphorylation and heme metabolism, as well as increased angiogenic signaling, consistent with heightened epithelial stress and metabolic reprogramming. In contrast, iEoE exhibited reduced activity of epithelial–mesenchymal transition (EMT) and myogenesis pathways, consistent with attenuated epithelial remodeling signatures given the disease control relative to aEoE at the time of sampling.

More granular Reactome pathway and curated gene set enrichment analysis revealed specific mechanistic modules underlying these molecular processes (Supplemental Fig. 1D, G; Table S7, S8). Compared to controls, EoE was associated with alterations in DNA methylation, RNA polymerase I promoter opening, DNA replication pre-initiation, metallothionein-mediated metal binding, and RHO GTPase effector signaling. Immune-mediated pathways such as IL-4/IL-13, IL-7, and IL-10 signaling, broader IL signaling networks, neutrophil degranulation, complement regulation, and lymphoid–non-lymphoid immunoregulatory interactions were also significantly enriched in EoE compared to controls. Additionally, pathways involving ECM remodeling, cytoskeletal dynamics, cell–cell interaction signaling, and developmental cell lineage programs reflected profound disruptions in epithelial structure and tissue remodeling. Comparisons between aEoE and iEoE revealed further mechanistic differentiation. Active disease showed stronger enrichment for epithelial stress and proliferative pathways, including cellular senescence, keratinization, and cornified envelope formation, indicating heightened injury responses. It also showed increased GPCR signaling, ECM degradation, and junctional and adhesion signaling, consistent with progressive epithelial destruction and tissue turnover.

To confirm the robustness of these transcriptomic differences, we reanalyzed an independent dataset from Jacobse et al. using the same analytical pipeline^12^. The EoE-related gene signatures reported in the pediatric cohort in the reference study were reproduced in our dataset (Supplemental Fig. 1F), and 80.4% of DEGs from our cohort overlapped with those from the reference study (Supplemental Fig. 1H), supporting the generalizability and reproducibility of the observed transcriptional patterns.

### Esophageal transcriptomic profiles reflect EoE activity rather than clinical severity

After assessing the transcriptomic derangements associated with EoE activity, we examined whether the transcriptomic patterns aligned more closely with EoE activity, as defined by the PEC, or with EoE severity, as measured by the I-SEE scores. PCA involving all EoE samples revealed that children with high I-SEE scores, including three classified as aEoE and one as iEoE, with scores above 20, did not segregate into a distinct transcriptional cluster. Instead, they grouped with their respective EoE activity groups defined by PEC (Fig. 2A). These findings suggest that while I-SEE captures the broader clinical and phenotypic burden of the disease, the esophageal transcriptional profile primarily reflects local histologic inflammation. Consistent with this distinction, one patient in our cohort exhibited a disproportionately high I-SEE score, driven by malnutrition and endoscopic abnormalities, despite being classified as iEoE by PEC (Fig. 2B), underscoring the discordance between tissue eosinophilia and global disease severity described in the I-SEE framework.

**Fig. 2.**
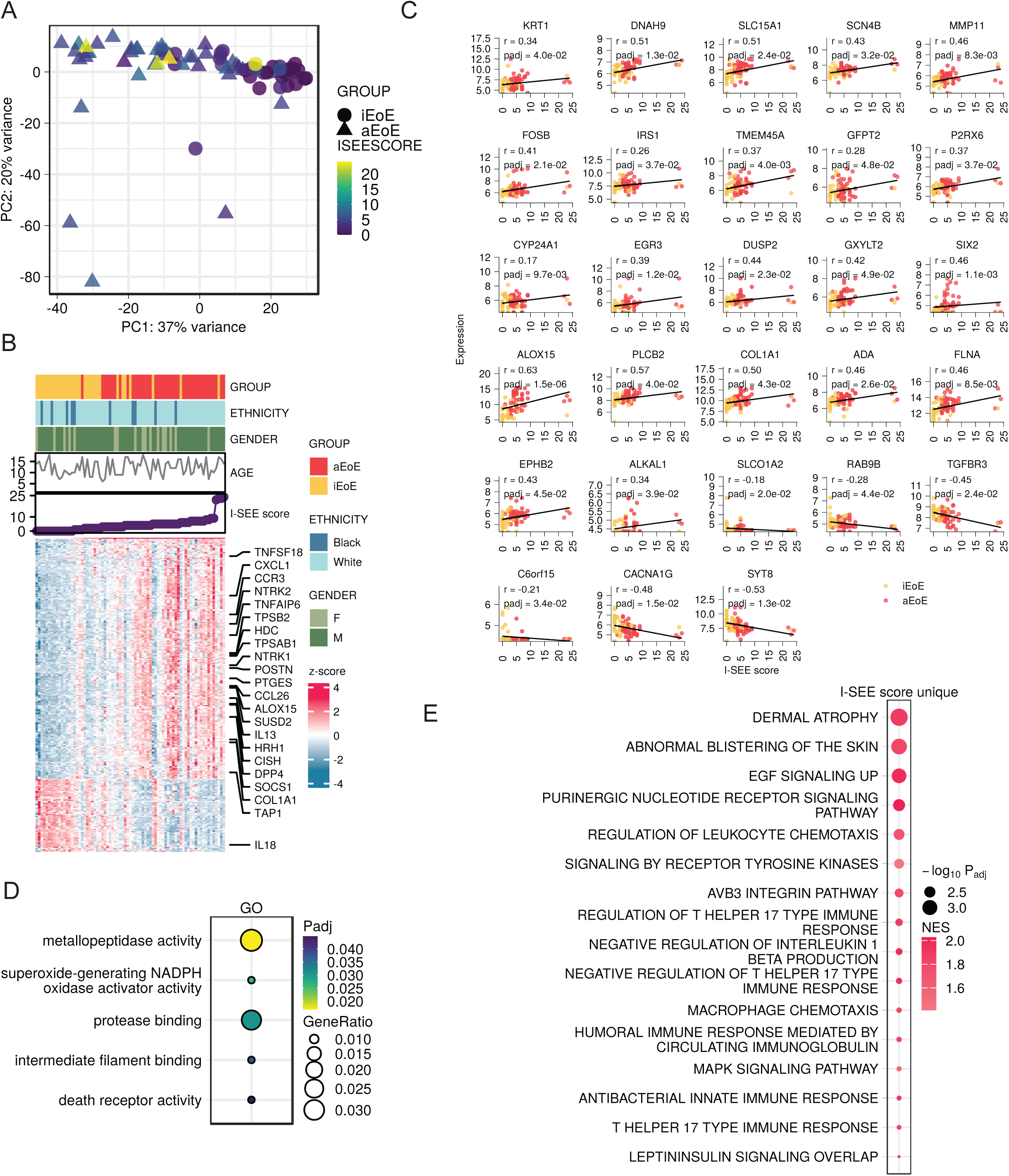
A) PCA plot of the whole transcriptome from all EoE patients. Circles are iEoE while triangles are aEoE. Colors represent I-SEE scores. B) Differential expression genes associated with the I-SEE score identified in the EoE group in our cohort. Data batch, ethnicity, gender, and age were adjusted as covariates. C) Scatter plots of 28 ISEE-score associated genes showing the relationship between gene expression and I-SEE score. The y-axis shows the expression values after normalization with the variance-stabilizing transformation function from DESeq2. Correlation coefficients of Spearman correlation tests were shown. Adjusted p values are from the DE analysis with covariates included. D) The significant GO terms only identified with the I-SEE score-associated genes. E) Significant Hallmark gene sets from I-SEE score-associated pre-ranked GSEA analysis.

### Genes associated with clinical severity highlight hallmark EoE pathobiology

Next, to identify the molecular features associated with disease severity, we correlated transcriptomic profiles with I-SEE scores. In all, 333 severity-associated genes were identified (Fig. 2B; Table S5). A majority of them overlapped with DEGs that distinguished aEoE from controls and exhibited moderate-to-strong correlations with I-SEE score (Fig. 2B) and encompassed hallmark pathways of EoE pathogenesis, including epithelial remodeling, barrier dysfunction, and immune activation. There were 28 I-SEE–specific genes converging on epithelial injury–response and maladaptive remodeling programs rather than canonical type 2 inflammation, highlighting pathways involved in cytoskeletal reorganization and mechanotransduction (FLNA, KRT1, EPHB2), extracellular matrix remodeling and fibrostenotic signaling (COL1A1, MMP11, TGFBR3), and stress-adaptive metabolic and oxidative responses (ALOX15, GFPT2, CYP24A1) (Fig. 2C). Together, they point to epithelial structural instability, altered cell–matrix interactions, and metabolic reprogramming as key molecular features of EoE severity that are distinct from eosinophil-driven disease activity.

To identify gene ontologies uniquely associated with I-SEE scores, we filtered enrichment terms shared with any group-level comparisons of EoE activity. Five GO terms remained unique to I-SEE–related genes (Fig. 2D), implicating epithelial cell death, cytoskeletal destabilization, proteolytic activity, oxidative stress, and extracellular matrix turnover. Complementary GSEA revealed additional I-SEE-associated programs (Fig. 2E), including T-helper 17 differentiation, leukocyte chemotaxis, cytokine regulatory networks, extracellular matrix organization, integrin signaling, and cell–cell adhesion pathways, as well as EGFR/MAPK signaling and metabolic stress responses. Collectively, these findings indicate that increasing severity of EoE reflects not only intensified type 2 inflammation but also broader activation of epithelial injury responses and progressive fibrostenotic remodeling, indicating a mechanistic shift toward tissue-level dysfunction.

### Severity of EoE is associated with impaired epithelial differentiation and stromal-immune activation

To assess how the severity-associated transcriptional programs were related to the cellular-level changes, we deconvolved cell-type proportions using BayesPrism with a previously published matched human esophageal scRNA-seq reference dataset (Supplementary Fig. 2A-D). The inferred compositions aligned with the single-cell estimates (Supplementary Fig. 2E), supporting robust mapping of cell-type abundance across disease states.

Analysis of individual epithelial subsets revealed distinct shifts across EoE activity (Fig. 3A). Controls and iEoE samples showed higher proportions of quiescent, Trans1, and differentiated epithelial cells. In contrast, aEoE samples were enriched for proliferating epithelial cells, consistent with prior scRNA-seq findings^6^, suggesting impaired maturation and increased epithelial turnover in active disease. In addition to the epithelial changes, mast cells, endothelial cells, and fibroblasts were expanded in aEoE relative to iEoE and controls, aligning with their roles in inflammation, angiogenesis, and extracellular matrix remodeling.

**Fig. 3.**
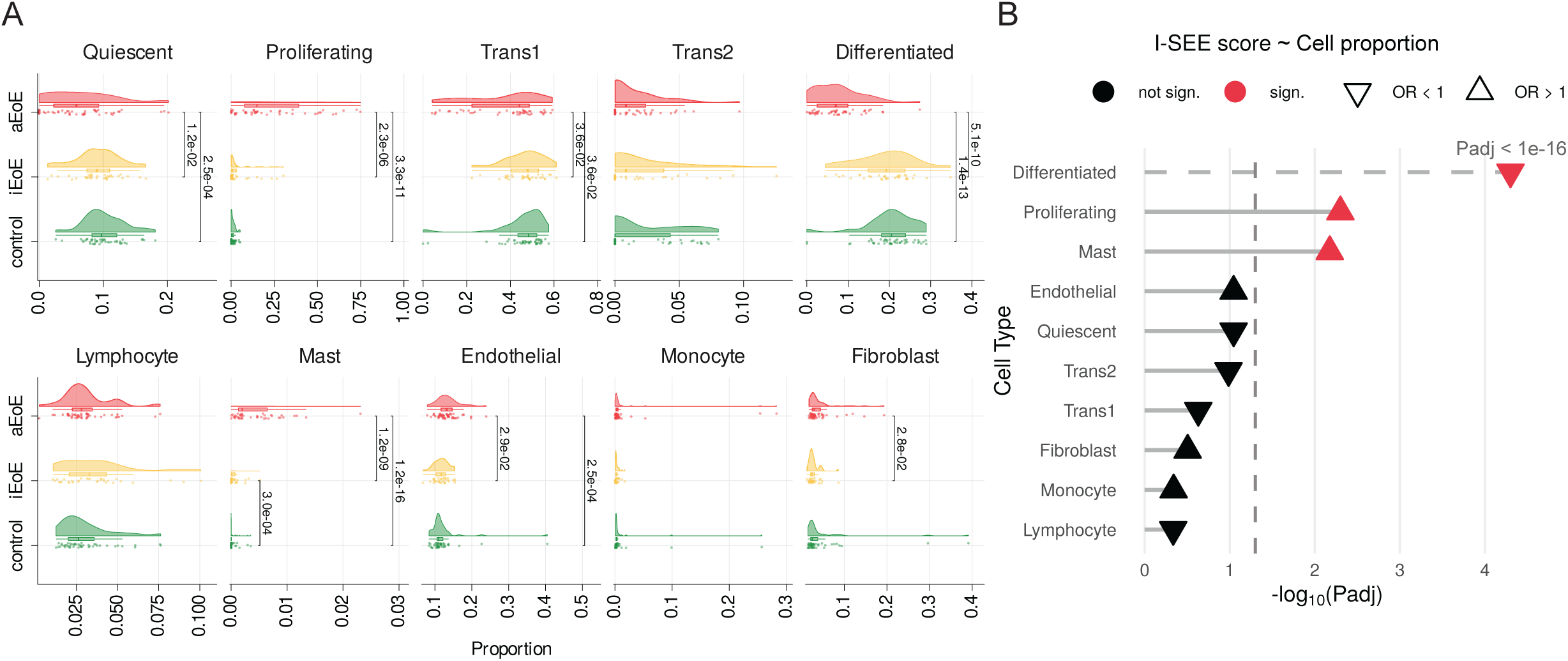
A) The raincloud plots for each cell type in our cohort. Cell proportions were compared between sample groups for each cell type using the Wilcox test. Multiple test adjusting was performed with BH procedure. B) Lollipop plot showing association between I-SEE score and cell types. Each lollipop represents a different cell type, with the horizontal bar length proportional to −log₁₀(adjusted p-value) and the endpoint shape indicating the direction of association. Red endpoints denote statistically significant associations (adjusted p-value < 0.05), while black endpoints denote non-significant associations. Triangle-up shapes (Δ) indicate positive associations (odds ratio > 1), while triangle-down shapes (∇) indicate negative associations (odds ratio < 1). The dashed vertical line marks the significance threshold (adjusted p-value = 0.05). The dashed lollipop stick for the “Differentiated” cell type indicates an extremely low adjusted p-value (labeled as “Padj < 1e-16”). Cell types are ordered vertically by statistical significance.

To directly link these cellular changes with EoE severity, we modeled the association between cell-type proportions and I-SEE scores using multivariate linear regression. Notably, proliferating epithelial cells and mast cells were positively associated with I-SEE scores. In contrast, differentiated epithelial cells showed a significant inverse relationship (Fig. 3B). The inferred proportions of mast cells were consistently low across samples, reflecting their relative scarcity within esophageal biopsies. As a result, estimates of mast cell–severity associations are inherently more sensitive to small proportional fluctuations and may be less robust than those derived from abundant epithelial populations. In contrast, epithelial subsets accounted for the vast majority of cellular content and exhibited coordinated, severity-linked shifts across proliferative, transitional, and differentiated states. These findings suggest that while mast cells may act as potent amplifiers of inflammatory and remodeling signals, the clinical severity captured by I-SEE is more plausibly driven by widespread epithelial state dysfunction rather than by changes in rare immune populations alone. These results indicate that increasing EoE severity, as assessed by I-SEE scores, reflects a shift toward epithelial immaturity and mast cell-driven inflammation, supported by intensification of stromal activation and tissue remodeling programs.

### Proliferating and transitional epithelial states are associated with EoE severity

To determine whether specific epithelial states contribute disproportionately to EoE severity, we applied Scissor^13^ by integrating bulk RNA-seq data annotated with I-SEE scores and the matched esophageal scRNA-seq reference dataset. Scissor identifies individual single cells whose transcriptional profiles are most strongly associated with the phenotype of interest, leveraging shared gene-expression correlations to link patient-level disease severity to specific cellular states^13^. It has previously been used to highlight the roles of immune-evasion-related subpopulations and molecular mechanisms in breast cancer^14^, and in our study, it allowed us to precisely identify the cell populations most responsible for driving EoE severity.

Using the I-SEE score for severity, Scissor identified 1,677 positively associated (Scissor-positive) and 2,158 negatively associated (Scissor-negative) cells (Fig. 4A). Strikingly, nearly 100% phenotype-associated Scissor-positive and Scissor-negative cells belonged to epithelial lineages (Fig. 4B), suggesting that epithelial cellular dysfunction, rather than immune cell variation, is the dominant cellular correlate of EoE severity.

**Fig. 4.**
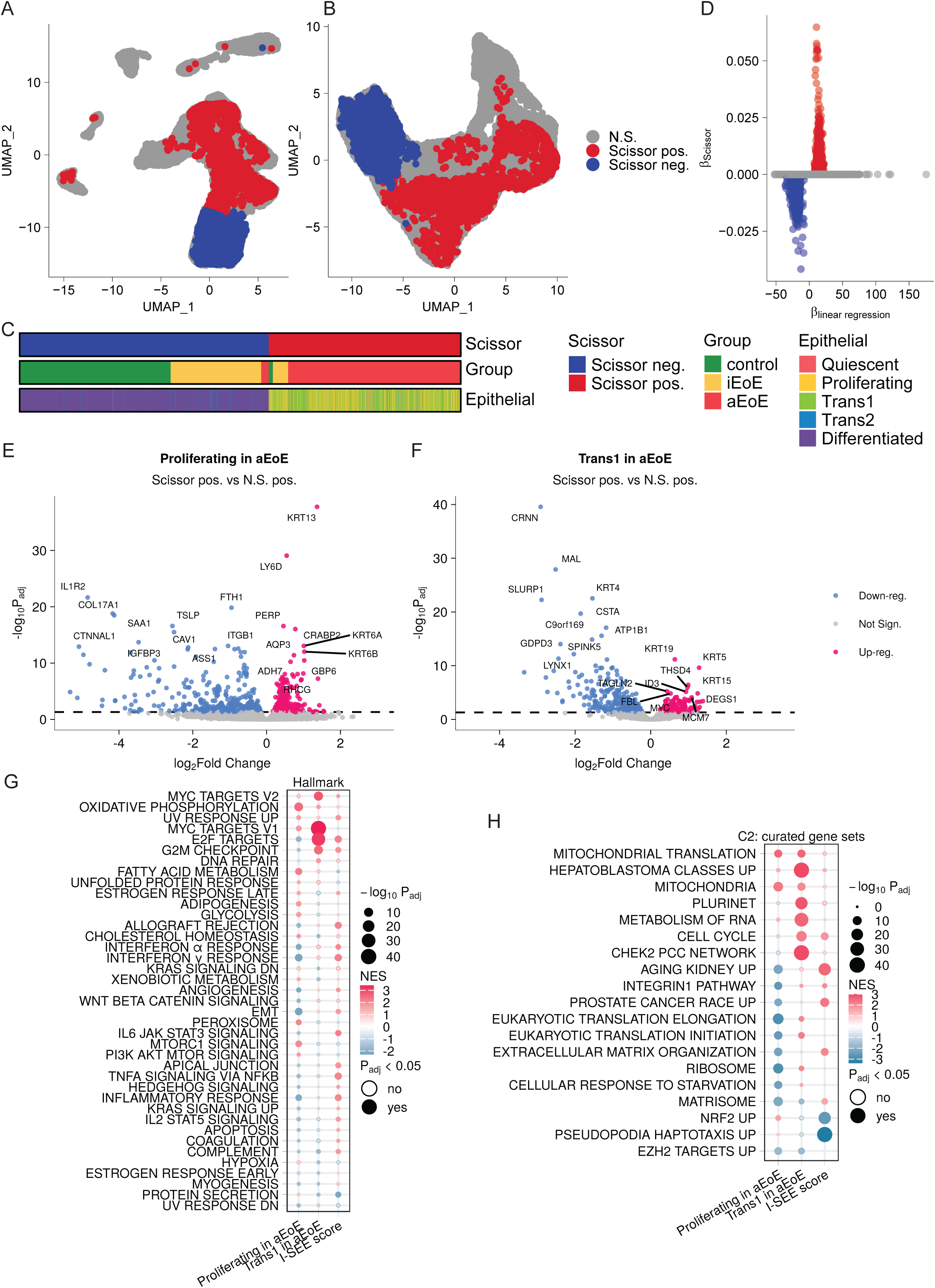
Distribution of the Scissor positive and Scissor negative cells identified by Scissor in scRNAseq data (Rochman et al., 2022) on a UMAP plot of: A) all cells, B) epithelial cells, C) EoE activity groups and epithelial cell types. Cells were sorted by Scissor groups and EoE activity groups. D) Scatter plot of coefficients of the simple linear regression model and Scissor model of correlation matrix of single cell and bulk sample with I-SEE score. Volcano plot of DE genes between Scissor positive and Scissor negative cells in E) proliferating epithelial cells in aEoE patients, F) Trans1 epithelial cells in aEoE patients. G) Significant Hallmark gene sets from pre-ranked GSEA analysis of scissor pos. vs non-significantly positive (N.S. pos.) in epithelial cell subtypes in aEoE patients. H) Significant curated gene sets from pre-ranked GSEA analysis of Scissor positive vs non-significantly positive in epithelial cell subtypes in aEoE patients. Redundant gene sets within curated and Hallmark categories were excluded.

With the cellular origins of EoE severity-associated signatures defined, we next examined how these populations vary by EoE activity and epithelial subtype. We found that Scissor-positive cells were derived mainly from aEoE samples (89.5 percent), whereas only 3 percent of Scissor-negative cells originated from aEoE. At the level of epithelial subtypes, Scissor-positive cells were enriched for proliferating epithelial cells (34.2%) and Trans1 transitional epithelial cells (61.3%), representing immature or remodeling states associated with the higher I-SEE scores (Fig. 4C). In contrast, 97.3% of Scissor-negative cells were differentiated epithelial cells, reflecting a mature and functionally intact epithelium corresponding to lower I-SEE scores.

### Defining the spectrum of epithelial cells based on the intensity of association with EoE severity

To contextualize the biology of Scissor-positive epithelial cells, we examined how these cells differ from other epithelial cells of the same subtype and EoE activity group. By integrating the linear regression coefficients, reflecting the association between I-SEE scores and the cell–bulk correlation matrix, with the Scissor-derived coefficients (Fig. 4D), we stratified epithelial cells into six groups according to their strength of association with the I-SEE score (Supplemental Fig. 3A). The details of the group definition are provided in the Methods. Notably, most of the Scissor-positive cells localized to proliferating (29.6 percent) and Trans1 (55 percent) epithelial lineages from aEoE, reinforcing the central role of epithelial expansion and transitional remodeling in driving disease severity in pediatric EoE (Supplemental Fig. 3B). These positive severity-linked cells were observed across patients and not restricted to a single specimen, indicating that they do not reflect subject-specific artifacts (Supplemental Fig. 3C).

### Distinct pathway programs distinguish proliferating versus transitional severity-associated cells

As the Scissor-positive and Scissor-negative cells are inherently linked to EoE activity status and epithelial lineage, we performed activity- and subtype-matched comparisons within proliferating and Trans1 epithelial populations from aEoE samples to isolate I-SEE associated transcriptional programs.

Within proliferating epithelial cells of aEoE samples, Scissor-positive cells exhibited transcriptional signatures distinct from non-significantly positive proliferating cells. Scissor-positive cells showed upregulation of KRT13, a marker of early epithelial differentiation, and downregulation of COL17A1 and IL1R2, genes involved in epithelial adhesion and immunoregulatory function (Fig. 4E). Meanwhile, within the Trans1 compartment, Scissor-positive cells showed marked suppression of mature epithelial markers compared with non-significantly positive Trans1 cells, including CRNN, MAL, and KRT4, which are characteristic of terminal squamous differentiation (Fig. 4F). Distinct DEG sets were identified with these epithelial subtype-specific Scissor-positive cells (Supplementary Fig. 3D).

Functional enrichment analyses against GO Biological Process and Reactome pathway database show that, Scissor-positive proliferating epithelial cells were characterized by a coordinated activation of biosynthetic and metabolic machinery, including cytoplasmic translation, peptide-chain elongation, viral mRNA translation, selenocysteine synthesis, and GCN2-mediated amino-acid–deficiency responses, alongside structural remodeling pathways such as desmosome organization, epithelial regeneration, and extracellular matrix organization (Supplementary Fig. 3E, F). These signatures indicate heightened protein synthesis, metabolic stress adaptation, and regenerative remodeling. In contrast, Trans1 epithelial cells showed pronounced dedifferentiation and barrier destabilization, characterized by loss of mature epithelial markers and enrichment of pathways involved in keratinization, epidermal differentiation, chemotaxis, leukocyte recruitment, cytokine-mediated signaling, and regulation of peptidase activity (Supplementary Fig. 3E, F).

Investigation through pre-ranked GSEA against Hallmark and curated gene sets further distinguished the two epithelial states (Fig. 4G, H). Severity-associated proliferating epithelial cells showed enrichment of oxidative phosphorylation, MTORC1 signaling, PI3K AKT MTOR signaling, and fatty acid metabolism, reflecting elevated metabolic and biosynthetic demands supporting epithelial turnover under inflammatory stress. In contrast, Scissor-positive Trans1 epithelial cells exhibited enrichment of cell-cycle-associated programs, including E2F targets, G2M checkpoint, MYC targets, cell cycle, and DNA repair, indicating heightened proliferative signaling within a dedifferentiated transitional state linked to higher I-SEE scores. Together, these findings delineate two complementary epithelial mechanisms underlying EoE severity. Proliferating epithelial cells engage a metabolically driven regenerative program characterized by elevated biosynthesis and structural remodeling, whereas transitional epithelial cells adopt an immune-interacting, dedifferentiated remodeling program marked by barrier destabilization and proliferative signaling. The coexistence of these programs suggests a coordinated epithelial response that integrates metabolic activation with immune-mediated remodeling, thereby amplifying tissue dysfunction and contributing to disease burden in active EoE.

### Single-cell epithelial programs reveal features not detectable in bulk RNA-seq

To evaluate how these epithelial programs relate to bulk molecular signatures behind EoE severity, we compared Scissor-derived signatures with I-SEE-associated bulk RNA-seq pathways. These DEGs derived from epithelial subsets show minimal overlap with the DEGs associated with I-SEE score identified in our bulk RNAseq data (Supplementary Fig. 3D). Bulk signatures were dominated by IL responses, inflammatory signaling pathways, and metabolic suppression, reflecting signals from diverse mixed cell types (Fig. 2D, E). In contrast, the single-cell analysis uncovered epithelial-intrinsic injury programs, including loss of differentiation, translational activation, oxidative stress responses, and epithelial junctional remodeling (Fig. 4G, H; Supplementary Fig. 3E, F). These epithelial changes were not apparent in bulk RNA-seq analyses, where immune-derived inflammatory signals predominate, but were revealed by single-cell approaches that resolve epithelial injury and remodeling programs.

Some genes involved in canonical EoE pathways were also modulated, including CD74, HLA-DRA, HLA-DRB1 (antigen-presenting pathways), S100A9 (acute inflammatory response), IFITM3 (interferon responses), IVL, SPRR3, TGM3, TSLP (epithelial stress and remodeling programs), and POSTN (Type 2 inflammatory signaling). They are hallmarks of chronic mucosal inflammatory diseases such as EoE (Supplemental Fig. 3G). Collectively, these analyses define severity-associated transcriptional programs within proliferating and transitional epithelial subtypes and identify the epithelial cell states most strongly correlated with increasing I-SEE clinical severity.

## Discussion

EoE is a chronic and progressive immunoinflammatory disease. In children, in addition to esophageal symptoms of feeding difficulties, vomiting, and abdominal pain, it can result in extraesophageal complications such as growth faltering, nutritional deficiencies, and impaired quality of life. I-SEE score captures the multidimensional burden of EoE by integrating clinical, endoscopic, histologic, and complications into a composite score that can be used in clinical practice. To systematically elucidate the pediatric-specific molecular and cellular correlates of this onerous disease, we leveraged the largest single-center pediatric EoE cohort and compared transcriptomic features across both EoE activity groups and I-SEE scores using integrated bulk RNA sequencing and single-cell–informed computational analyses.

Bulk transcriptomic profiling revealed that aEoE displayed a distinct molecular profile characterized by widespread immune activation, epithelial stress, and extracellular matrix remodeling (Fig. 1A-F). In contrast, transcriptomic patterns in iEoE closely resembled those of controls, with minimal dysregulation. We also noted a stepwise gradient across control, iEoE, and aEoE states reflected in gene-level signatures spanning type 2 inflammation, mast-cell activation, epithelial barrier structure, neuro-immune signaling, innate immune activity, and tissue remodeling (Fig. 1D). Pathway-level analyses further revealed coordinated disturbances in IL signaling, chromatin regulation, ECM turnover, cytoskeletal organization, and metabolic programs, delineating both shared and severity-specific processes (Fig. 1E-F; Supplemental Fig. 1D, G). Upon comparing pediatric EoE transcriptomic data with a recent EoE tissue transcriptome meta-analysis integrating multiple independent cohorts across both pediatric and adult populations^12^, we observed strong concordance between the two datasets.

These signatures were independently replicated, supporting the robustness and generalizability of our findings (Supplemental Fig. 1F, H). Incorporating the I-SEE clinical severity framework provided additional biological insight beyond classifying EoE activity. EoE severity correlated with genes that recapitulate canonical EoE biology and with pathways uniquely associated with clinical severity, including epithelial cell death, cytoskeletal destabilization, oxidative stress responses, and extracellular matrix remodeling (Fig. 2B-E). These findings reflect a combination of inflammatory activation, epithelial injury, and structural remodeling, consistent with the multidomain nature of the I-SEE score. Together, these findings suggest that increasing I-SEE severity reflects coordinated changes in inflammatory pathways and epithelial adaptation, highlighting dynamic crosstalk between immune activation and epithelial remodeling.”

A significant innovation of this study was the use of Scissor to link clinical severity to single-cell epithelial states. Scissor identified epithelial subpopulations whose expression profiles were most strongly associated with I-SEE scores (Fig. 4A-C). Nearly all severity-associated cells were epithelial rather than immune-derived (Fig. 4B), indicating that epithelial biology plays a central role in shaping clinical severity. Two epithelial subpopulations, proliferating epithelial cells and Trans1 epithelial cells, accounted for the majority of severity-associated cells. Although both were strongly associated with I-SEE scores, they exhibited distinct biological programs. Scissor-positive proliferating cells showed enrichment for translational, metabolic, and regenerative remodeling programs, including cytoplasmic translation, peptide-chain elongation, amino-acid metabolic pathways, and desmosome and extracellular matrix organization, together indicating heightened protein synthesis, metabolic stress adaptation, and immature regenerative activity (Supplementary Fig. 3E, F; Fig. 4G, H). In contrast, Scissor-positive Trans1 cells were enriched for epithelial dedifferentiation and immune-interacting pathways, including keratinization, cytokine-mediated signaling, chemotaxis, and junctional remodeling, alongside proliferative hallmarks (Supplementary Fig. 3E, F; Fig. 4G, H), consistent with barrier destabilization and active tissue restructuring. Gene-level analyses further highlighted upregulation of KRT13 and suppression of adhesion-related and differentiation markers (COL17A1, IL1R2, CRNN, MAL, KRT4) in severity-associated cells (Fig. 4E-F).

These epithelial programs were obscured mainly in bulk RNA-seq, which was dominated by inflammatory signals from mixed cell types. The single-cell–resolved findings revealed epithelial-intrinsic adaptations, including translational activation, oxidative stress responses, junctional restructuring, and loss of differentiation programs and features that define epithelial plasticity in severe disease but are not detectable at the tissue level (Fig. 4E-H; Fig. 2E). Cell-type deconvolution further supported this framework: proliferating epithelial cells and mast cells increased with higher I-SEE scores, while differentiated epithelial cells declined (Fig. 3A-B), linking epithelial immaturity and stromal-immune activation with clinical severity.

Together, these findings support a model in which severe EoE reflects not only amplified inflammation but also a profound shift in epithelial composition and function. Expansion of proliferating and transitional epithelial states, accompanied by depletion of mature epithelial cells, indicates a disrupted differentiation trajectory and persistent epithelial injury. These cellular states offer insight into the cellular programs associated with the symptomatic, endoscopic, and histologic features captured by the I-SEE score, including edema, fragility, fibrosis, and stricturing.

This study has limitations. Although this study utilized the largest single-center pediatric EoE cohort and validated transcriptomic and single-cell signatures in an independent dataset, broader evaluation with differing racial distributions and clinical management strategies will be essential to further establish generalizability. The cross-sectional study design limits our ability to infer temporal dynamics of the epithelial severity state and how it evolves across different treatment modalities, or to predict long-term outcomes such as fibrosis or stricture formation. We had relatively few patients in the severe I-SEE category or with fibrostenotic complications. This is consistent with our current understanding that children typically present with an inflammatory phenotype and over time, with suboptimal control of eosinophilic inflammation, they may develop fibrostenotic complications in adolescence or young adulthood. Computational phenotype linking approaches, such as Scissor, while powerful, depend on model assumptions that warrant further validation in prospective single-cell studies. In addition, transcriptomic profiling was limited to esophageal mucosal biopsies; therefore, molecular changes in deeper layers of the esophageal wall were not assessed. As a result, some clinical manifestations that contribute to I-SEE scores reflecting deeper tissue involvement may not have been captured by mucosal sampling alone. Finally, esophageal tissue profiling does not capture extraesophageal manifestations of EoE, which may also influence severity scores.

Despite these limitations, this study has several strengths. First, transcriptome signatures of severity-associated epithelial cells identified through Scissor might serve as cell-state-specific biomarkers for disease monitoring or predicting therapy response. Second, therapeutic strategies aimed at restoring epithelial differentiation, stabilizing transitional epithelial populations, or mitigating epithelial metabolic and oxidative stress may complement cytokine-targeting agents to achieve deeper disease control.

Finally, integrating clinical scoring with bulk and single-cell transcriptomics provides a framework for understanding disease severity in other chronic inflammatory disorders characterized by epithelial dysfunction.

In summary, this study identifies the molecular and cellular determinants of clinical severity in pediatric EoE. By integrating clinical phenotyping with bulk transcriptomics and single-cell–resolved analysis, we demonstrate that epithelial remodeling, loss of differentiation, and altered metabolic and structural states are central features of severe disease. These translational insights provide a foundation for precision severity stratification and therapeutic approaches aimed at targeting the epithelial drivers of EoE progression.

## Methods

### Sample collection

Prospectively collected distal esophageal biopsies for research and corresponding clinical metadata under Vanderbilt Institutional Review Board-approved protocols (# 151341 and 160785) for previously published studies were analyzed^15, 16^. Briefly, children aged 6-18 years with a prior diagnosis of EoE or upper gastrointestinal symptoms suggestive of EoE and undergoing an esophagogastroduodenoscopy (EGD) at Vanderbilt Children’s Hospital were enrolled. During the EGD, 2-3 biopsies, each from the proximal and distal esophagus (4-6 per participant), were collected for clinical care and submitted for histopathologic assessment per our institutional protocol. In addition, two distal esophageal biopsies (≤ 5 cm from the lower esophageal sphincter) were collected for research purposes and stored in RNA later (Qiagen, catalog number 76154) separately at −80°C. Children with known esophageal injury or surgery, celiac disease, neurodevelopmental disorders, enteral tube-fed patients, inflammatory bowel disease, and exposure to antibiotics and systemic steroids within the past 30 days were excluded to minimize confounding.

### Clinical data and activity indices

Demographic (age at EGD, sex, ethnicity), clinical (weight, indication for EGD, allergic co-morbidities, medication exposure including exposure to PPI and TS), endoscopic (esophageal mucosal abnormalities visualized during EGD and rated per the validated endoscopic reference score [EREFS]^17^, and histologic (number of esophageal biopsies, PEC, and EoEHSS^18^ information was gathered from the electronic medical records. I-SEE score to quantify EoE severity was calculated across three domains: symptoms and complications, inflammatory features, and fibrostenotic features. Each feature was assigned a point value (1-15), and the total score was used to classify the disease into inactive (0), mild (1-6 points), moderate (7-14 points), and severe (≥ 15 points). In our analysis, we treated I-SEE scores as a continuous severity measure to preserve the full range of clinical variation and to improve statistical power for correlation with transcriptomic features. Notably, specific features, such as esophageal perforation, malnutrition with body mass index < the 5^th^ percentile, or a decreased growth trajectory, automatically assigned a patient to the severe category due to their clinical significance^19^.

### RNA extraction and metatranscriptomic sequencing of esophageal biopsies

Total RNA was extracted from research esophageal biopsy specimens that had been flash-frozen and stored at –80 °C, following previously described protocols^15, 20^. Illumina sequencing libraries were prepared using the NEBNext Ultra II RNA Library Prep Kit (NEB #E7775). Library quality and fragment size distribution were assessed using the Agilent Bioanalyzer High Sensitivity RNA chip. Libraries were sequenced on the Illumina NovaSeq X Series platform, generating 2 × 150 bp paired-end reads with an average sequencing depth of approximately 40 million paired-end reads per sample, as described in our previous publication^15^.

### Bulk RNAseq analysis

Reads were trimmed to remove adapter sequences using Cutadapt (v4.8). Quality control on both raw reads and adaptor-trimmed reads was performed using FastQC (v0.12.1) (www.bioinformatics.babraham.ac.uk/projects/fastqc). Reads were aligned to the Gencode GRCh38.p13 genome using STAR (v2.7.11a)^21^. Gene annotations from Gencode v38 were provided to STAR to improve mapping accuracy. FeatureCounts (v2.0.6)^22^ was used to count the number of mapped reads to each gene. For differential expression gene identification, DESeq2 was used with a linear regression model, including data batch, ethnicity, gender, and age as covariates. When extracting the result tables, lfcThreshold was set to 0.5 for most models, and FDR-adjusted p-value <= 0.05 was considered as significantly differentially expressed genes (v1.42.1)^23^. Function enrichment analysis was performed against Gene Ontology, KEGG, and Reactome pathway databases with differentially expressed genes using the ClusterProfiler package^24^. Pre-ranked GSEA analysis was performed using the fgsea r package against the MsigDB database (v2025.1.Hs).

### Cell type deconvolution analysis

scRNAseq data and processed Seurat objects were provided by Dr. Rothenberg’s group^7^. BayesPrism was used to perform cell type deconvolution with the reference scRNAseq dataset^25^. Cell compositions were compared between EoE activity groups with the Wilcoxon rank-sum test or fitted in the linear regression model for continuous clinical outcomes (i.e., I-SEE score).

### Integrative phenotype–single-cell analysis using Scissor

To determine which cell populations were associated with clinical disease severity, we used Scissor, a computational framework that integrates bulk RNA-seq phenotypes with single-cell gene-expression data^13^. Scissor is specifically designed to identify the individual cells within a single-cell dataset whose transcriptional profiles best reflect the clinical traits observed across bulk tissue samples by modeling the clinical outcomes with the bulk-single cell correlation matrix. In this study, our clinical phenotype of interest was the I-SEE score, a validated measure of EoE severity. We first generated a cell–sample similarity matrix by quantifying the transcriptional similarity between each single cell in the reference scRNA-seq dataset and each bulk RNA-seq sample from our cohort. Scissor then applied a regression model that uses this similarity matrix along with the I-SEE scores to determine which single cells exhibit gene-expression patterns that rise or fall with increasing disease severity.

Cells whose transcriptional profiles correlated positively with higher I-SEE scores were classified as Scissor-positive, indicating that these cells are associated with more severe EoE. Conversely, cells whose profiles correlated with lower scores were classified as Scissor-negative, representing features more consistent with remission or milder disease. Because Scissor shrinks coefficients of non-significant cells to zero, we further quantified each cell’s association with I-SEE scores using a linear regression model applied to the same cell–bulk correlation matrix. Some cells retained high regression coefficients despite having Scissor coefficients of zero (Fig. 4D). To better classify epithelial cells with minimum influence of different models, we defined six groups: (1) 1,677 Scissor-positive cells; (2) 2,158 Scissor-negative cells; (3) “potential Scissor positive” cells whose regression coefficients fell within mean ±3 SD of Scissor-positive cells; (4) “potential Scissor negative” cells whose coefficients fell within mean ±3 SD of Scissor-negative cells; (5) non-significantly positive cells (coefficients > 0 after exclusion of prior groups); and (6) non-significantly negative cells (coefficients < 0 after exclusion of prior groups). The distribution of regression coefficients and the number of cells in each category across epithelial subtypes and sample groups are shown in Supplemental Figures 3A and 3B. To ensure that signals were not driven by a small number of subjects, we evaluated Scissor-positive cell proportions across samples (Supplemental Figure 3C). This integrative strategy allowed us to pinpoint the specific immune and epithelial subpopulations that drive the transcriptomic signature of clinical severity, providing cellular-level insight that cannot be achieved with bulk RNA-seq or single-cell sequencing alone.

Scissor has been validated in cancer and other disease contexts^26–28^, and its application here offers a powerful means of identifying clinically meaningful cell states in pediatric EoE.

Pseudo-bulk differential expression analysis was performed between Scissor-positive and non-significantly positive cells in each subtype of epithelial cells in aEoE, rather than iEoE and control, due to the cell number limitation. The same strategies for DE identification, function enrichment, and GSEA analyses were adopted afterwards, as described in the bulk RNAseq analysis, to maintain consistency.

### Data availability

The bulk RNA-sequencing data generated in this study was deposited in the Gene Expression Omnibus (GEO) under accession number GSE313910.

De-identified clinical data supporting the findings of this study are available from the corresponding author upon reasonable request and pending appropriate institutional approvals.

### Code availability

All custom code used for preprocessing, statistical analysis, and visualization of the bulk and single-cell transcriptomic data has been deposited in a public GitHub repository and is accessible at https://github.com/YuWang-VUMC/EoE_severity_transcriptome.

## Acknowledgement

Y.C. is supported by R01DK141803.

M.E.R is supported by NIH R01 AI045898, R01 AI124355, R01 NS105715, U19AI070235, and the

Campaign Urging Research for Eosinophilic Disease (CURED) Foundation.

G.H. is supported by NIH grants K23DK131341 and R21AI168832.

S.V.R, Y.W and S.Q is supported by NIH grant R21AI168832.

The authors acknowledge Ms. Regina Tyree for managing the tissue biorepository and database.

The content is solely the responsibility of the authors and does not necessarily represent the official views of the NIH.

## Author Contributions

Y.W. contributed to the analysis and interpretation of data, drafting the manuscript, and critical revision of the manuscript.

P.G. contributed to bulk RNA sequencing.

M.K.S. contributed to the data analysis.

J.M.P. contributed to interpretation of data, and critical revision of the manuscript.

Y.A.C. contributed to the interpretation of data and the critical revision of the manuscript.

H-W.L. contributed to the analysis and interpretation of data and the critical revision of the manuscript.

M.E.R. contributed to the interpretation of data and the critical revision of the manuscript.

Q.S. contributed to the analysis and interpretation of data and made critical revisions to the manuscript.

G.H. and S.V.R. contributed to study conceptualization, data acquisition and analysis, manuscript drafting and critical revision, and securing funding. Each author has approved the final draft submitted.

## Supplementary Figures

Supplemental Fig. 1. A) PCA plot of the whole transcriptome from the entire cohort. EoE groups are represented by colors. B) Scatter plots of log2FC between EoE vs. control and aEoE vs. control. Significant DE genes are shown in colors. Correlation tests were performed, and the correlation coefficients and p-values are shown in the plots. C) Scatter plots of log2FC between aEoE vs. control and aEoE vs. iEoE. Significant DE genes are shown in colors. Correlation tests were performed, and the correlation coefficients and p-values are shown in the plots. D) Top 10 significant Reactome pathways of each comparison from function enrichment analysis. Redundant pathways were filtered. E) Differential expression genes identified by comparing aEoE versus iEoE. Data batch, ethnicity, gender, and age were adjusted as covariates. Red indicates up-regulated in aEoE, while blue indicates up-regulated in iEoE. F) Differential expression genes identified by comparing EoE versus control in pediatric patients from the Jacobse et al. cohort. Data batch, gender, and age were adjusted as covariates. EoE-related genes were shown in the plots. Red indicates up-regulation in EoE, while blue indicates up-regulation in control. G) Significant curated gene sets encompassing at least one comparison from pre-ranked GSEA analysis. Color indicates the normalized enrichment score, while point size indicates the adjusted p-value (-log_10_Padj). Up-regulated pathways in the numerator group are shown in red, while the down-regulated pathways in the numerator group are shown in blue. H) Venn diagram of DE genes from 3 comparisons: EoE vs control, aEoE vs iEoE from this study, and EoE vs control in pediatric patients in the Jacobse et al. I) I-SEE score distributions across duration time of the disease. Kruskal-Wallis test was applied to check the differences between the categorized duration of EoE.

Supplemental Fig. 2. A) UMAP plot of the reference scRNAseq data for all major cell types. UMAP plot of the reference scRNAseq data for B) all major cell types by sample groups, C) scRNAseq data for all epithelial subtypes, D) scRNAseq data for all epithelial subtypes grouped by sample groups. E) Cell type composition in our cohort and the scRNAseq reference dataset. The box with solid outlines shows the cell-type proportions in our cohort, while the box with dashed outlines shows the cell-type proportions in the scRNAseq reference data. Colors indicate different EoE activity groups for the samples.

Supplementary Fig. 3. A) Coefficient distribution of defined cell groups across epithelial subtypes and sample groups from the linear regression model. Six cell groups were defined: non-significantly negative (N.S. neg.), potential Scissor-negative (poten. neg.), Scissor-negative, non-significantly positive (N.S. pos.), potential Scissor-positive (poten. pos.), and Scissor-positive cells. B) Stacked bar plot of cell counts of defined cell groups across epithelial subtypes and sample groups. C) Scissor-positive cell proportions in each epithelial subtype across 5 aEoE samples. D) Venn diagram of DE genes from 3 analyses: scissor pos. vs. N. S. pos. proliferating epithelial cells from aEoE group; scissor pos. vs. N. S. pos. Trans1 epithelial cells from aEoE group; genes associated with I-SEE score from bulk RNAseq data of our cohort. E) Top 10 significant GO Biological Process terms from function enrichment analysis between Scissor-positive versus non-significantly positive groups in proliferating and Trans 1 epithelial cells from aEoE. Corresponding results related to the I-SEE score from bulk RNAseq were included for comparison. Redundant terms were filtered with simplify function (cutoff = 0.6, by = “p.adjust”, select_fun = min, measure = “Wang”). F) Top 10 significant Reactome pathways from function enrichment analysis between Scissor-positive versus non-significant positive groups in proliferating and Trans 1 epithelial cells from aEoE. Corresponding results related to the I-SEE score from bulk RNAseq were included for comparison. Redundant terms were filtered with simplify function (cutoff = 0.6, by = “p.adjust”, select_fun = min, measure = “Wang”). G) Canonical EoE pathway related DEGs between Scissor-positive versus non-significantly positive groups in epithelial subtypes.

## Table Legends

Table S1. Clinical characteristics of the cohort in this study

Table S2. Significantly different expression genes between aEoE vs control

Table S3. Significantly different expression genes between EoE vs control

Table S4. Significantly different expression genes between aEoE vs iEoE

Table S5. Significantly different expression genes associated with the I-SEE score

Table S6. Statistically significant functional enrichment GO items of DE genes from 4 comparisons: aEoE vs control, EoE vs control, aEoE vs iEoE, and I-SEE score

Table S7. Statistically significant functional enrichment KEGG pathways of DE genes from 4 comparisons: aEoE vs control, EoE vs control, aEoE vs iEoE, and I-SEE score

Table S8. Statistically significant functional enrichment Reactome pathways of DE genes from 4 comparisons: aEoE vs control, EoE vs control, aEoE vs iEoE, and I-SEE score

Table S9. Statistically significant gene sets identified by preranked GSEA based on aEoE vs control

Table S10. Statistically significant gene sets identified by preranked GSEA based on EoE vs control

Table S11. Statistically significant gene sets identified by preranked GSEA based on aEoE vs iEoE

Table S12. Statistically significant gene sets identified by preranked GSEA based on the I-SEE score

Table S13. Statistically significant different expression genes between scissor pos. vs n. s. pos. in Proliferating epithelial cells in active EoE group

Table S13. Statistically significant different expression genes between scissor pos. vs n. s. pos. in Trans1 epithelial cells in active EoE group

Table S15. Statistically significant functional enrichment GO items of DE genes from 2 comparisons: scissor pos. vs n. s. pos. in Proliferating and Trans1 epithelial cells in active EoE group

Table S16. Statistically significant functional enrichment Reactome pathways of DE genes from 2 comparisons: scissor pos. vs n. s. pos. in Proliferating and Trans1 epithelial cells in active EoE group

Table S17. Statistically significant gene sets identified by preranked GSEA based on the gene ranks of scissor pos. vs n. s. pos. in Proliferating epithelial cells in active EoE group

Table S18. Statistically significant gene sets identified by preranked GSEA based on the gene ranks of scissor pos. vs n. s. pos. in Trans1 epithelial cells in active EoE group.

