## Supplementary material for "Esophageal epithelial cell-state transitions underlie the severity of pediatric eosinophilic esophagitis": Table S2

| **Table S1. Clinical characteristics of the cohort** | | | |  |  |  |
| --- | --- | --- | --- | --- | --- | --- |
|  |  | **control (N=46)** | **iEoE (N=32)** | **aEoE (N=43)** | **Total (N=121)** | **p value** |
| **Age (Years)**^@^ |  | 13 (7 - 17) | 11 (7 - 18) | 12 (6 - 18) | 12 (6 - 18) | 0.75^2^ |
| **Ethnicity**^#^ | White | 43 (93%) | 27 (84%) | 38 (88%) | 108 (89%) | 0.43^1^ |
|  | Black | 3 (7%) | 5 (16%) | 5 (12%) | 13 (11%) |  |
| **Sex**^#^ | Male | 16 (35%) | 26 (81%) | 32 (74%) | 74 (61%) | < 0.01^1^ |
| **Batch** | batch10914 | 5 (11%) | 8 (25%) | 5 (12%) | 18 (15%) | 0.37^1^ |
|  | batch10956 | 20 (43%) | 8 (25%) | 15 (35%) | 43 (36%) |  |
|  | batch7191 | 16 (35%) | 13 (41%) | 15 (35%) | 44 (36%) |  |
|  | batch4996 | 5 (11%) | 3 (9%) | 8 (19%) | 16 (13%) |  |
| **Disease Duration**^#^ | < 6 months | 22 (50%) | 4 (12%) | 4 (9%) | 30 (25%) | < 0.01^1^ |
|  | 6 -12 months | 14 (32%) | 0 (0%) | 5 (12%) | 19 (16%) |  |
|  | 13 - 24 months | 4 (9%) | 6 (19%) | 8 (19%) | 18 (15%) |  |
|  | > 24 months | 4 (9%) | 22 (69%) | 26 (60%) | 52 (44%) |  |
| **History of**^#^ | Esophageal dilation | 0 (0%) | 0 (0%) | 4 (9%) | 4 (3%) | 0.02^1^ |
|  | Food impaction | 1 (2%) | 4 (12%) | 10 (23%) | 15 (12%) | 0.01^1^ |
| **Allergic comorbidities**^#^ | Food allergy | 8 (17%) | 10 (31%) | 21 (49%) | 39 (32%) | < 0.01^1^ |
|  | Seasonal allergy | 5 (11%) | 21 (66%) | 16 (37%) | 42 (35%) | < 0.01^1^ |
|  | Allergic rhinitis | 8 (17%) | 20 (62%) | 23 (53%) | 51 (42%) | < 0.01^1^ |
|  | Eczema | 7 (15%) | 16 (50%) | 11 (26%) | 34 (28%) | < 0.01^1^ |
|  | Asthma | 9 (20%) | 16 (50%) | 16 (37%) | 41 (34%) | 0.02^1^ |
| **Medication exposure**^#^ | Antihistamines | 0 (0%) | 2 (11%) | 1 (5%) | 3 (7%) | 0.89^1^ |
|  | Nasal TS | 0 (0%) | 0 (0%) | 2 (10%) | 2 (5%) |  |
|  | PPI | 1 (100%) | 15 (79%) | 15 (71%) | 31 (76%) |  |
|  | TS | 0 (0%) | 0 (0%) | 1 (5%) | 1 (2%) |  |
|  | PPI + TS | 0 (0%) | 2 (11%) | 2 (10%) | 4 (10%) |  |
| **Clinical symptoms** | Known EoE | 1 (2%) | 32 (100%) | 36 (84%) | 69 (57%) | < 0.01^1^ |
|  | Abdominal pain | 33 (72%) | 0 (0%) | 2 (5%) | 35 (29%) |  |
|  | Difficulty swallowing | 5 (11%) | 0 (0%) | 4 (9%) | 9 (7%) |  |
|  | Vomiting | 3 (7%) | 0 (0%) | 0 (0%) | 3 (2%) |  |
|  | Nausea | 0 (0%) | 0 (0%) | 1 (2%) | 1 (1%) |  |
|  | Others | 4 (9%) | 0 (0%) | 0 (0%) | 4 (3%) |  |
| **Endoscopy** |  |  |  |  |  |  |
| Features^#^ | Edema | 1 (2%) | 4 (12%) | 27 (63%) | 32 (26%) | < 0.01^1^ |
|  | Exudates | 0 (0%) | 0 (0%) | 1 (2%) | 1 (1%) |  |
|  | Furrows | 0 (0%) | 3 (9%) | 4 (9%) | 7 (6%) |  |
|  | Rings | 1 (2%) | 0 (0%) | 0 (0%) | 1 (1%) |  |
|  | Stricture | 0 (0%) | 0 (0%) | 1 (2%) | 1 (1%) |  |
| Total EREFS^@^ |  | 0 (0.0 - 2.0) | 0 (0.0 - 2.0) | 2 (0.0 - 5.0) | 0 (0.0 - 5.0) | < 0.01^2^ |
| Phenotype^#^ | Inflammatory | 1 (2%) | 7 (22%) | 32 (74%) | 40 (33%) | < 0.01^1^ |
|  | Fibrostenotic | 1 (2%) | 1 (3%) | 10 (23%) | 12 (10%) | < 0.01^1^ |
| **Histology** | PEC^@^ |  | 2 (0 - 15) | 50 (18 - 112) | 2 (0 - 112) | < 0.01^2^ |
|  | Total EoEHSS^@^ |  | 0 (0 - 0.3) | 0.4 (0.1 - 1) | 0.2 (0 - 1) | < 0.01^2^ |
| **Severity** | I-SEE score^@^ |  | 1 (0 - 23) | 5 (1 - 24) | 4 (0 - 24) |  |
|  | I-SEE Categories^#^ |  |  |  |  |  |
|  | active |  | 15 (47%) | 0 (0%) | 15 (20%) |  |
|  | mild |  | 16 (50%) | 30 (70%) | 46 (61%) |  |
|  | moderate |  | 0 (0%) | 10 (23%) | 10 (13%) |  |
|  | severe |  | 1 (3%) | 3 (7%) | 4 (5%) |  |
| ^1^. Pearson’s Chi-squared test | |  |  |  |  |  |
| ^2^. Linear Model ANOVA | |  |  |  |  |  |
| ^@^ median (min-max); ^#^n (%); PPI: Proton-Pump Inhibitors; TS: Topical Steroids; PEC: Peak eosinophil count | | | | | |  |
