## Supplementary figures and images for "Esophageal epithelial cell-state transitions underlie the severity of pediatric eosinophilic esophagitis"

### Supplementary Figure 1

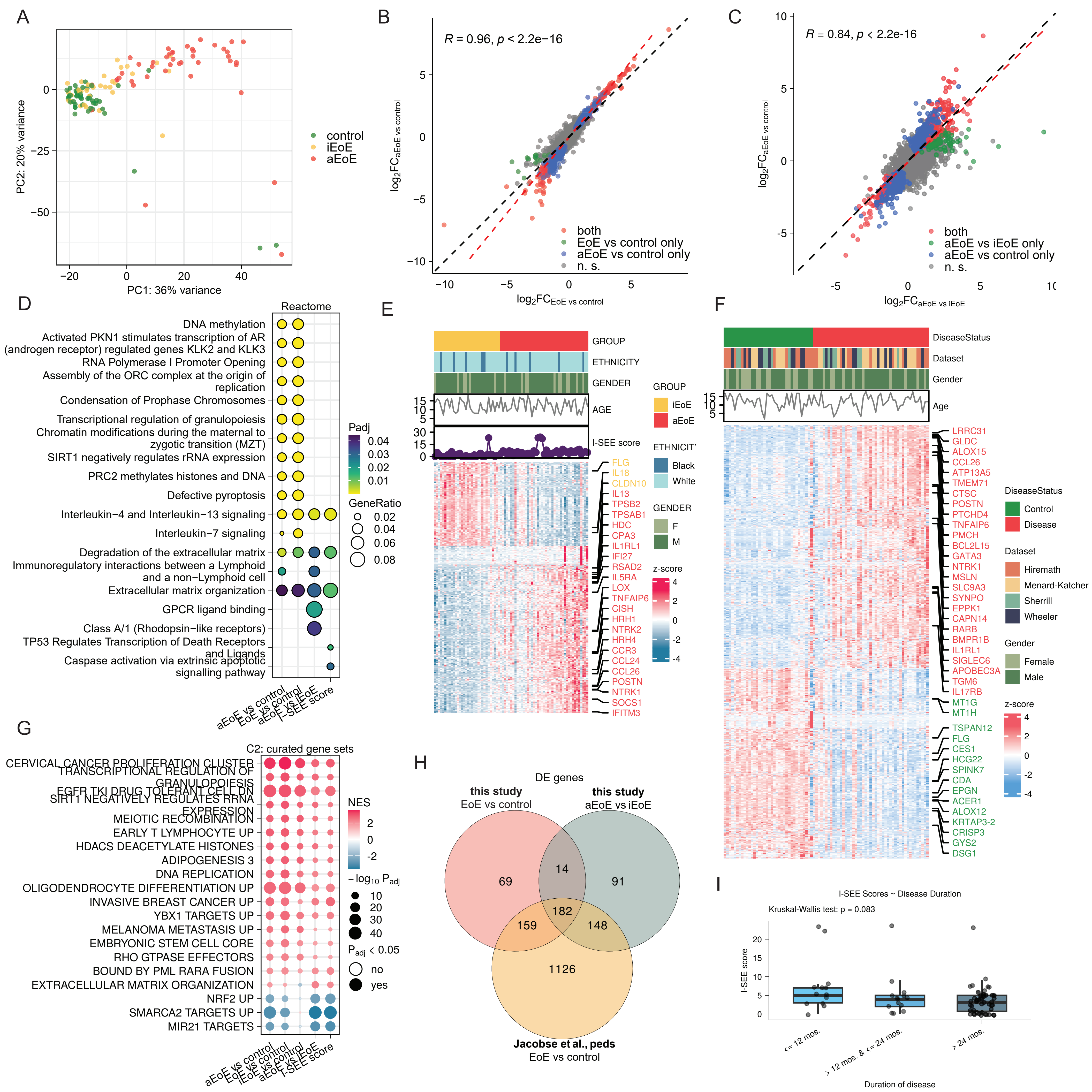

### Supplementary Figure 2

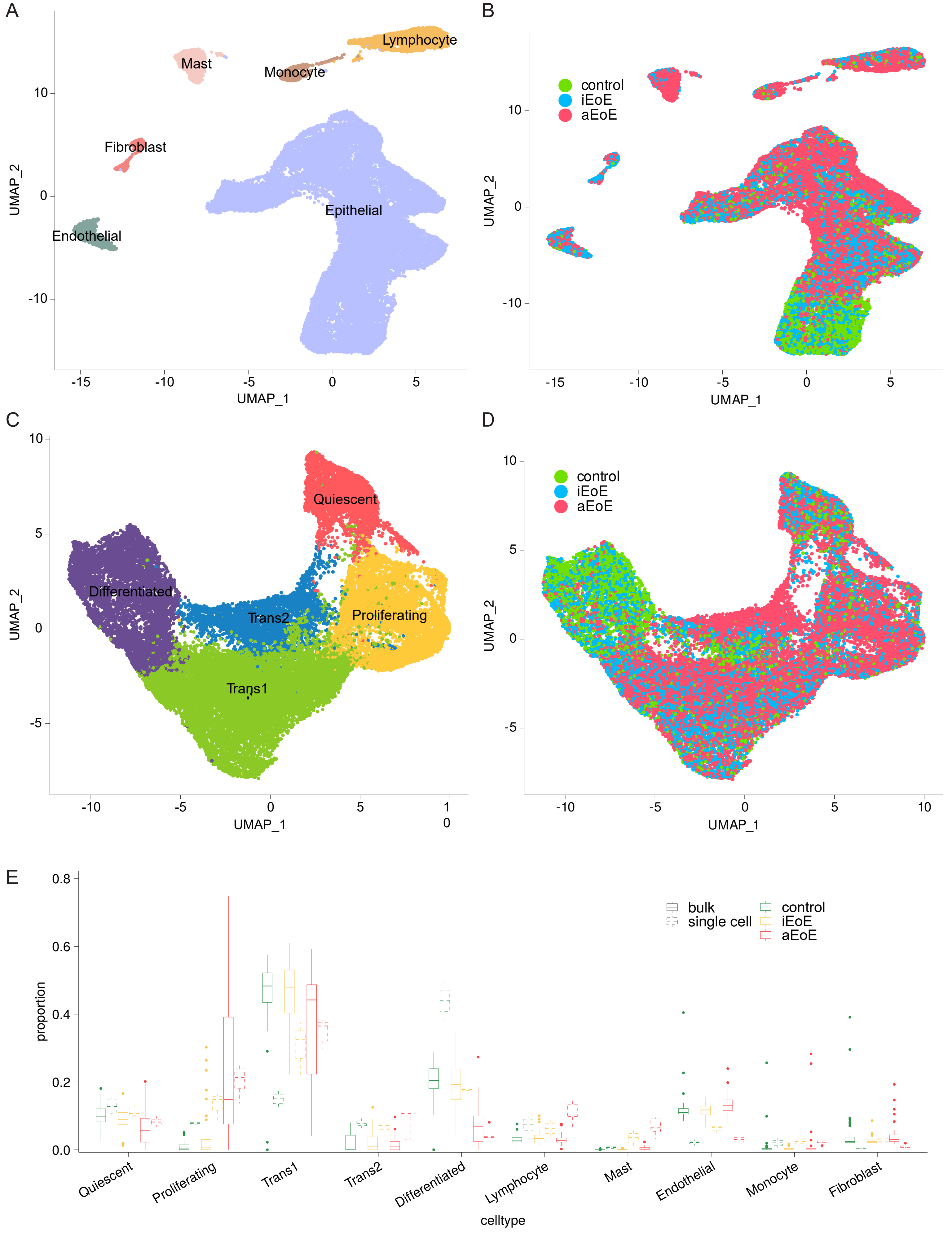

### Supplementary Figure 3

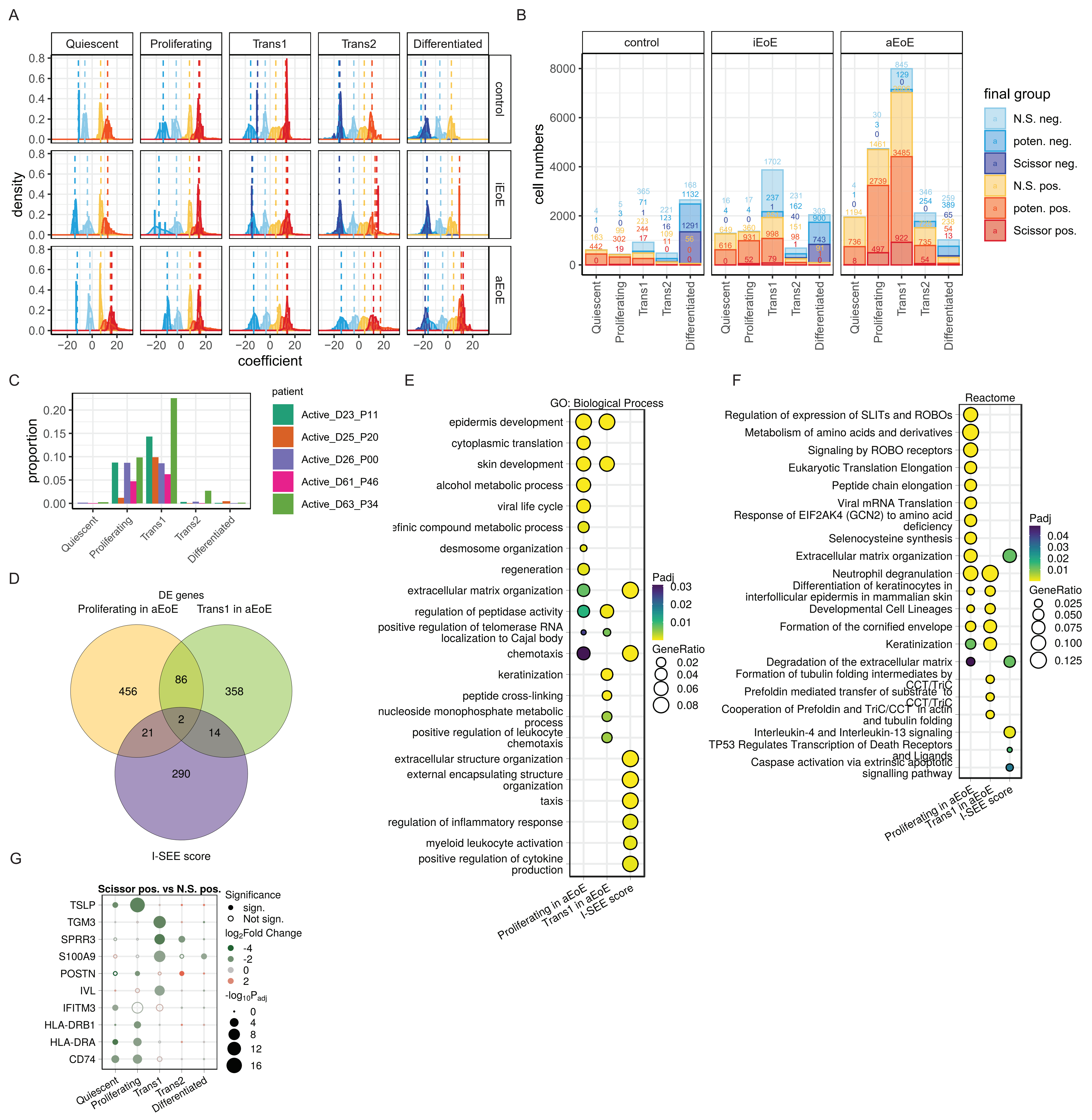
